# Resuscitation-promoting factor (Rpf) terminates dormancy among diverse soil bacteria

**DOI:** 10.1101/2024.11.10.622857

**Authors:** Jay T. Lennon, Brent K. Lehmkuhl, Lingling Chen, Melissa Illingworth, Venus Kuo, Mario E. Muscarella

## Abstract

Microorganisms often inhabit environments that are suboptimal for growth and reproduction. To survive when challenged by such conditions, individuals may engage in dormancy where they enter a metabolically inactive state. For this persistence strategy to confer an evolutionary advantage, microorganisms must be able to resuscitate and reproduce when conditions improve. Among bacteria in the phylum Actinomycetota, dormancy can be terminated by resuscitation-promoting factor (Rpf), an exoenzyme that hydrolyzes glycosidic bonds in the peptidoglycan of the cell wall. We characterized Rpf from *Micrococcus* KBS0714, a bacterium isolated from agricultural soil. Compared to previous studies, the Rpf elicited activity at relatively high concentrations, yet demonstrated high substrate affinity. Site-directed mutations at conserved catalytic sites significantly reduced or abolished resuscitation, as did the deletion of repeating motifs in a lectin-encoding linker region. We then tested the effects of recombinant Rpf from *Micrococcus* KBS0714 on a diverse set of dormant soil bacteria. Patterns of resuscitation mapped onto strain phylogeny, which reflected core features of the cell envelope. Additionally, the direction and magnitude of the Rpf effect were associated with functional traits, in particular, aspects of the moisture niche and biofilm production, which are critical for understanding persistence and resuscitation during dormancy. These findings expand our understanding of how Rpf may affect seed-bank dynamics and have implications for the diversity and functioning of soil ecosystems.

## 1 Introduction

Soils are among the most diverse microbial habitats on Earth [1, 2]. A single gram of soil can harbor thousands of co-occurring taxa ([3], and there is substantial turnover in composition across spatial scales [4]. One life-history strategy that may contribute to soil biodiversity is dormancy, a process that allows individuals to enter a reversible state of reduced metabolic activity [5]. It is estimated that less than 1% of the soil microbial biomass pool is metabolically active [6]. The remaining viable biomass is a ‘seed bank’. This reservoir of dormant individuals plays a crucial role in the evolution and persistence of populations, the maintenance of biodiversity, and the overall functioning of ecosystems [7–9].

Dormancy has independently evolved many times, giving rise to a variety of mechanisms by which microorganisms enter and exit metabolic states. The initiation of dormancy is often a response to environmental conditions that are unfavorable for growth and reproduction. For instance, some filamentous cyanobacteria form resting structures called akinetes in response to light or phosphorus limitation [10]. Fungal spore production is initiated in part by hormonal signaling between arbuscular mycorrhizae and the plant host [11]. Additionally, some bacteria enter dormancy by forming persister cells, which can be generated stochastically or in response to adverse environmental conditions [12].

Resuscitation is also crucial for the success of dormancy as a survival strategy. Since awakening at an inopportune time can be maladaptive, microorganisms often rely on environmental cues to time their resuscitation. This decision-making process is typically regulated by the detection of nutrients through membrane-bound receptors [13] or, in some cases, by more sophisticated mechanisms, such as tracking electrochemical potentials, a form of microbial memory that can be used to gauge environmental quality [14]. Another example comes from the Actinomycetota, a phylum of bacteria that emerge from dormancy using an extracellular enzyme known as resuscitation-promoting factor (Rpf). The ‘scout hypothesis’ proposes that these bacteria may stochastically awaken and secrete Rpf into the environment, thereby triggering the resuscitation of neighboring cells [15].

Resuscitation-promoting factors (Rpfs) were first identified in populations of *Micrococcus luteus*. When late stationary-phase cultures were exposed to filter-sterilized supernatants from actively growing cells, dormant bacteria rapidly resumed their metabolic activity [16]. Subsequently, the growth-stimulating compound was identified as a small (*∼* 16 kDa) protein that exhibits muralytic activity [17]. Rpf cleaves *β*-(1,4) glycosidic bond in peptidoglycan, a major constituent of the cell wall in nearly all bacteria [18, 19]. In addition to facilitating the remodeling and turnover of dormant cell walls necessary for outgrowth, the muropeptides released during Rpf-mediated hydrolysis may act as specialized signaling molecules that awaken closely related bacteria [20].

Rpf is broadly distributed among diverse lineages of the Actinomycetota [21, 22]. It is clinically relevant because Rpf can terminate latency in *Mycobacterium* strains, including those responsible for tuberculosis [23]. Rpf is also found among environmental isolates [24] and may be particularly important in soils. Metagenomic analyses indicated that up to 20% of all soil genomes contain Rpf homologs [5]. Meanwhile, experimental evidence suggests that Rpf has direct and indirect effects on seed bank dynamics through its influence on microbial diversity, soil respiration, and plant performance [25]. Very few studies, however, have characterized Rpf from soil or evaluated how it affects diverse strains of bacteria.

In this study, we conducted experiments with recombinant Rpf generated from an actinomycetotal strain, *Micrococcus* KBS0714, that was isolated from agricultural soil [26, 27]. After quantifying enzyme kinetics and concentration dependent responses, we characterized the effect of Rpf on the resuscitation dynamics of a diverse set of soil bacteria to determine the spectrum of activity in relation to phylogeny along with functional traits that are involved in soil dormancy.

## 2 Results

### 2.1 Rpf is a high affinity enzyme

We quantified the enzyme kinetics of purified recombinant Rpf from *Micrococcus* KBS0714 (Fig. S1) using fluorescein-labeled peptidoglycan (Fig. 1). When hydrolyzed, the fluorescent substrate products can be used to quantify muramidase activity. We characterized Rpf activity using the Michaelis-Menten function (see eq. 1). With maximum likelihood, we estimated that *V*_max_ was 59 units of fluorescein min^−1^ and that the *K*_*m*_ was 1.8 mg of peptidoglycan mL min^−1^. The relatively low *K*_*m*_ is consistent with reports of Rpf having high affinity and catalyzing reactions at low substrate concentrations [17].

**Fig 1.**
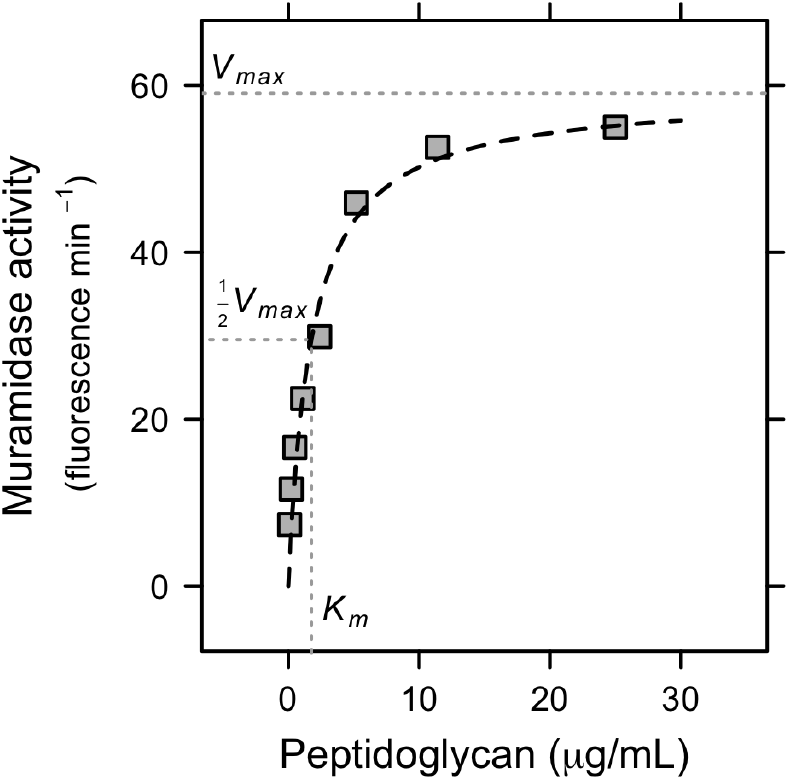
Enzyme kinetics of Rpf. We measured muramidase activity based on fluorescence released after purified Rpf (34 µM) was incubated with different concentrations of fluorescein-infused peptidoglycan. We fit the data to the Michaelis-Menten function (see eq. 1).

### 2.2 Rpf effect on growth is concentration-dependent

To evaluate the stimulatory or inhibitory effects of Rpf, we measured the biomass of *Micrococcus* KBS0714 following the incubation of dormant cells in R2 medium with different concentrations of recombinant protein. We fit the resulting data with the Monod growth model, which mathematically is identical to that of the Michaelis-Menten equation (see eq. 1). We determined that growth monotonically increased with Rpf concentration (Fig. 2). The half-saturation constant (*K*_*s*_) was 2.1 *µ*M Rpf and the estimated maximum biomass (OD500) was 2.3 (Fig. 2). Based on this relationship, there was no evidence for growth inhibition of *Micrococcus* KBS0714 at the elevated Rpf concentrations tested in our study. See Fig. S2 for discussion of negative controls.

**Fig 2.**
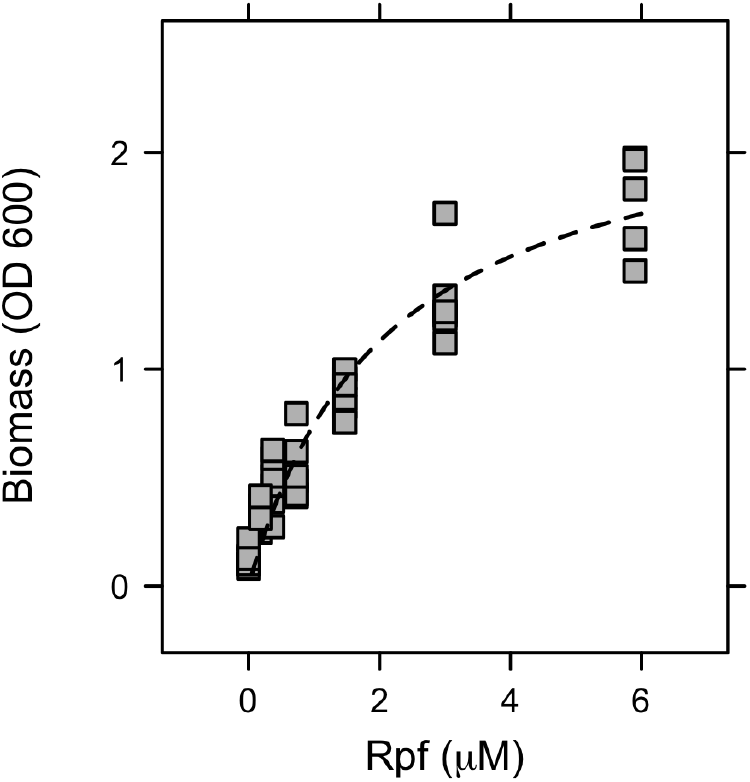
Concentration dependence of Rpf on bacterial growth. The regrowth of dormant *Micrococcus* KBS0714 in R2 medium with different concentrations of recombinant Rpf. We fit the data to the Monod function.

### 2.3 Rpf mutations interfered with resuscitation

When dormant cells were exposed to recombinant protein from the native *rpf* sequence, we observed a two-to four-fold increase in biomass (Fig. 3, *P*_adj_ *≤* 0.0005). This effect was diminished by site-directed mutagenesis of the conserved catalytic site (E54) within the *rpf* gene (see Fig. S3). When glutamic acid (E), a charged amino acid, was mutated to alanine (A), a hydrophobic amino acid, resuscitation by the recombinant Rpf was completely abolished (Fig. 3A, *P*_adj_ *<* 0.0001). However, resuscitation was only reduced by 40% when glutamic acid was mutated to lysine (K), a charged amino acid (Fig. 3B, *P*_adj_ *<* 0.0001). Similarly, when the glutamic acid residue was mutated to a polar uncharged amino acid, glutamine (Q), resuscitation was reduced by 25% (*P*_adj_ = 0.0005), with biomass still three-fold greater than the -Rpf control (*P*_adj_ *<* 0.0001, Fig. 3C).

**Fig 3.**
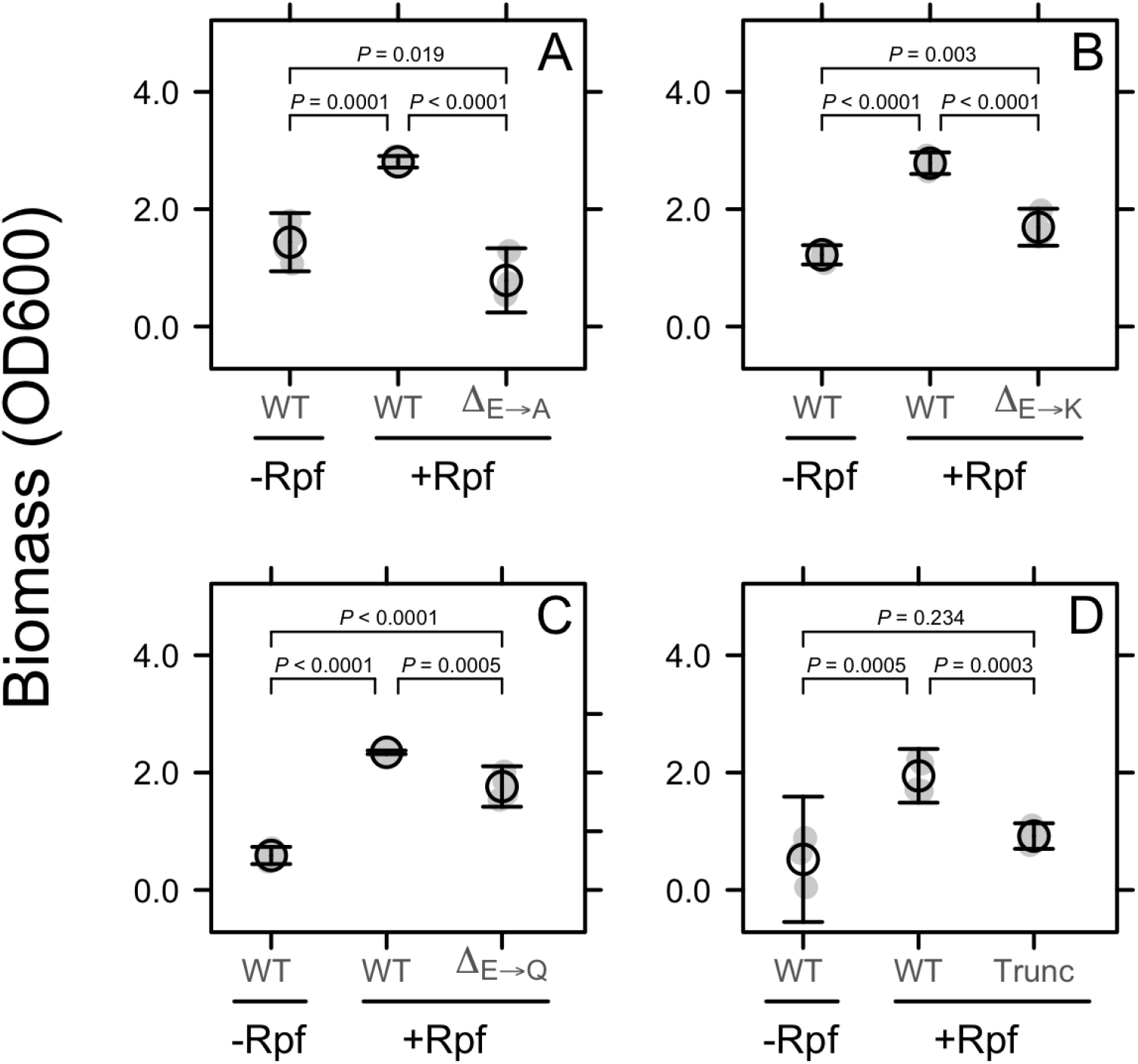
Rpf mutations reduced resuscitation. Effects of site-directed and truncation mutations on growth of *Micrococcus* KBS0714. Grey symbols represent raw data. Black symbols represent the mean ± SEM. In panels A–C, we used site-directed mutagenesis to change a conserved catalytic site (E54) from a glutamic acid residue (E) to an alanine (A), lysine (K), or glutamine (Q) residue. In panel D, we show results from a truncation mutation that deleted repeating motifs in a lectin-encoding linker region.

We identified structural variation in the amino acid sequence of *rpf* in (*Micrococcus* KBS0714). Compared to *Micrococcus luteus* (Flemming), which has been used as a model for studying Rpf [17], we identified a number of repeating motifs in a lectin-encoding linker region between the lysozyme-like domain and the LysM domain. In *Micrococcus* KBS0714, this linker region contains 21 tandemly arrayed repeats, whereas the Flemming strain contains only three repeats (Fig. S3). To assess whether this structural variation affected resuscitation, we deleted the variable linker region along with the LysM domain of Rpf. The activity of the resulting truncated Rpf was reduced by more than 50% (*P*_adj_ = 0.0003) and abolished resuscitation based on the fact that biomass was indistinguishable from the -Rpf control (*P*_adj_ = 0.234, Fig. 3D).

### 2.4 Rpf altered population dynamics

We quantified the effect of recombinant Rpf on the dynamics of a dormant population of *Micrococcus* KBS0714. Following 90 d of starvation, we transferred cells to lactate minimal medium. A one-time addition of Rpf (0.48 *µ*M) reduced lag time by 37% from 476 ± 27.1 h to 298 ± 3.4 h (*t*3.1 = 6.52, *P* = 0.007, Fig. 4). As a consequence, experimentally resuscitated populations entered exponential growth phase *∼* 7.5 d earlier than control populations. In contrast, Rpf had no effect on the maximum growth rate (*µ*max) (-Rpf = 0.058 ± 0.0213 d^−1^; +Rpf = 0.028 ± 0.0003 d^−1^; *t*_3.1_ = 1.40 *P* = 0.25) or the biomass yield (-Rpf = 4.1 ± 0.27; +Rpf = 3.6 ± 0.17, *t*_5.0_ = 1.68, *P* = 0.15) of *Micrococcus* KBS0714. However, Rpf minimized variabilitys among replicate populations in *µ*_max_ 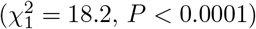and lag time 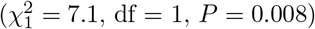 amounting to a 5-fold and 30-fold reduction, respectively, in the coefficient of variation (CV) of these growth-curve parameters.

**Fig 4.**
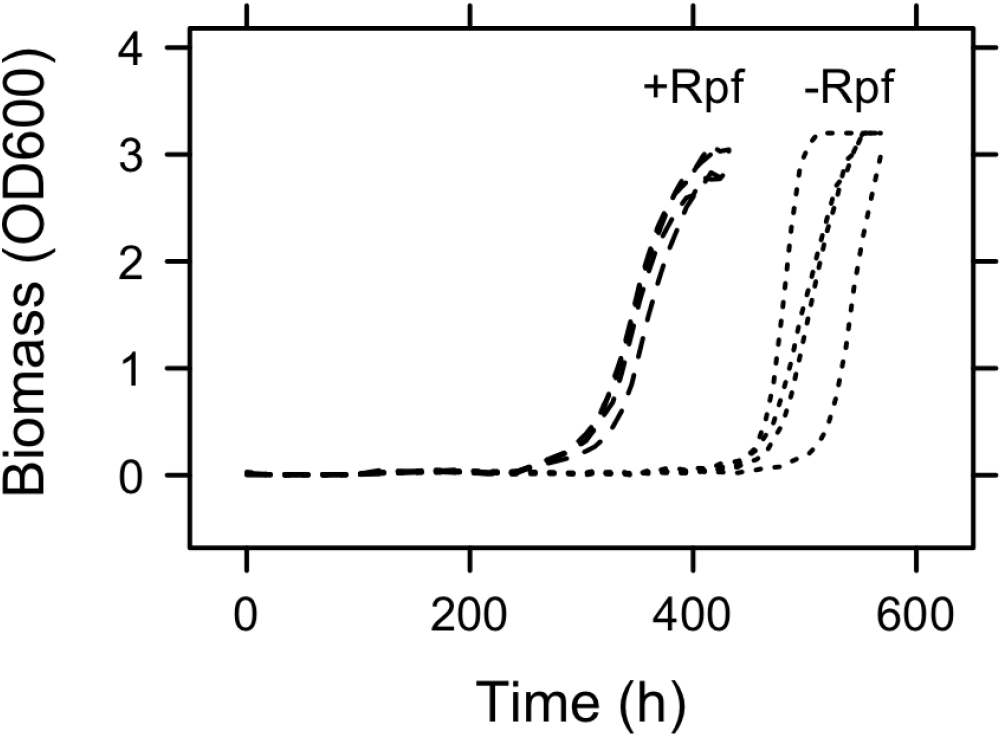
Population dynamics of dormant bacterial exposed to Rpf. Following 90 d of starvation, we transferred dormant *Micrococcus* KBS0714 into fresh medium with or without recombinant Rpf. The addition of Rpf reduced lag time by *∼*7.5 d. It also reduced among-population variability in *µ*_max_ and lag time indicating that Rpf may synchronize resuscitation-related processes.

### 2.5 Diverse bacteria affected by Rpf

To assess how diverse microorganisms resuscitate, we transferred dormant cells into fresh media with and without recombinant Rpf from *Micrococcus* KBS0714. During the resuscitation process, we quantified growth curve parameters (see eq. 2) and calculated the standardized Rpf effect size for each of the 12 strains of soil bacteria. There was a significant interaction between strain and Rpf treatment on growth yield (*F*_(1,11)_ = 3.35, *P* = 0009) and *µ*_max_ (*F*_(1,11)_ = 4.07, *P* = 0.0001) (Fig. S4). Some of this variation could be explained by the evolutionary history of the strains used in our study. For example, we detected phylogenetic signal for the Rpf effect size on *µ*_max_ (Pagel’s *λ*: 1.0, *P* = 0.013; Blomberg’s K: 0.99, *P* = 0.018) suggesting that aspects of resuscitation were consistent with a model of Brownian motion based on a maximum likelihood tree constructed from 16S rRNA gene sequences (Fig. 5).

**Fig 5.**
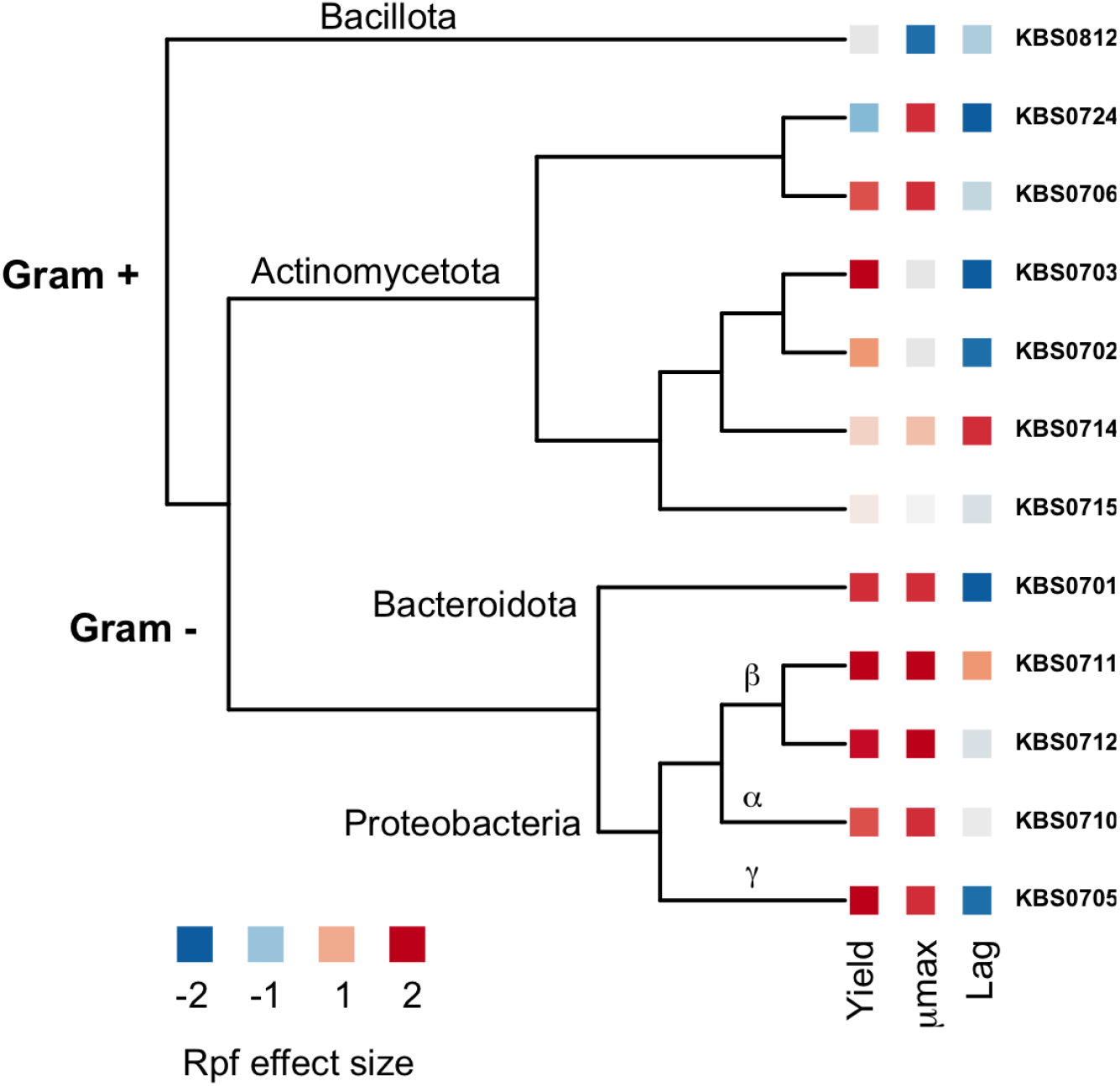
Effects of Rpf on the resuscitation of diverse soil bacteria from dormancy. Maximum likelihood tree based on 16S rRNA gene sequences from 12 strains of soil bacteria. For each strain of bacteria, we quantified growth curve parameters (i.e., *µ*_max_, yield, and lag time) in the presence (+Rpf) and absence (-Rpf) of recombinant protein (0.5 µmol/L). With that information, we calculated the Rpf effect size for each growth curve parameter.

In addition, variation in resuscitation among bacterial strains was associated with key features of the cell envelope. Specifically, Gram-positive strains responded differently to Rpf than Gram-negative strains. There was a significant interaction between these cell types and the Rpf treatment on growth yield (*F*_(1,82)_ = 9.47, *P* = 0.003) and *µ*_max_ (*F*_(1,82)_ = 7.30, *P* = 0.008). We found that Rpf had a 3.2-fold and 6.2-fold greater effect size on *µ*_max_ and growth yield, respectively, for Gram-negative bacteria compared to Gram-positive bacteria. In contrast, the Rpf effect size for lag time was similar for bacterial with contrasting cell types (Fig. 5).

Last, variation in the resuscitation of soil bacteria could be explained by functional traits. In particular, resuscitation was related to features of the moisture niche. We found that the Rpf effect size for both *µ*_max_ and growth yield increased linearly with the minimum water potential, a quantity that defines the water potential (MPa) where respiration (*R*) is equal to 5% of maximum respiration (*R*_*m*_*ax*). We also found that both *µ*_max_ and lag time decreased linearly with the breadth and optimum water potential of the bacterial moisture niche (Fig. 6). The Rpf effects size for lag time was not directly related to features of the moisture niche, but was positively correlated with biofilm production (*F*_(1,9)_ = 6.08, *P* = 0.036, *r*^2^ = 0.40), which can confer tolerance to desiccation stress.

**Fig 6.**
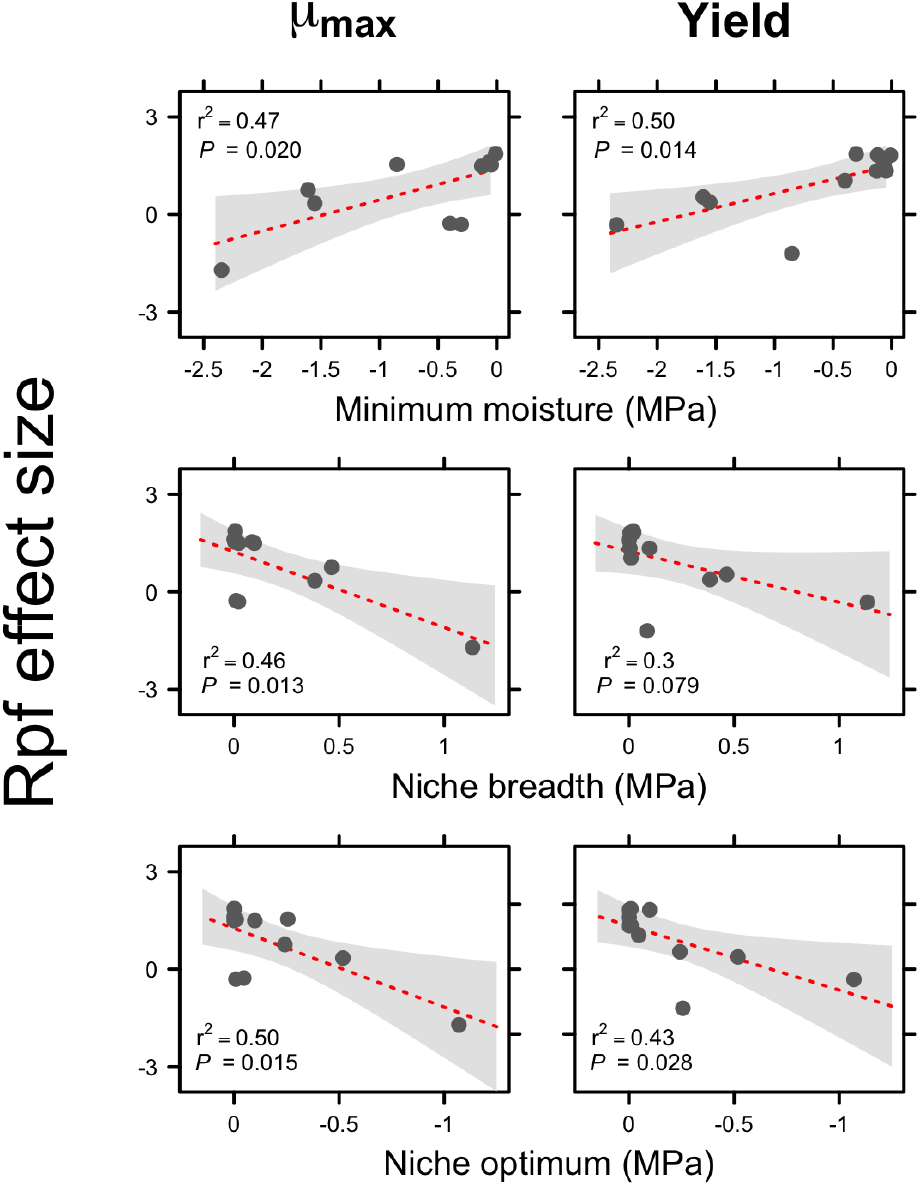
Relationship between Rpf effect size and functional traits. For each strain of bacteria, we transferred dormant bacteria to fresh medium in the presence (+Rpf) and absence (-Rpf) of recombinant protein (0.5 µmol/L). We then quantified resuscitation dynamics using parameters from the growth curve function (see eq. 2). With that information, we calculated the Rpf effect size for each of growth curve parameter (i.e., (*µ*_max_, yield, and lag time) and related it to different features of the moisture niche, which had been independently assessed [26], namely the minimum water potential, the niche breadth, and the niche optimum.

## 3 Discussion

Dormancy is a widespread survival strategy among microorganisms enabling them to persist in harsh and fluctuating environments. It has independently evolved across diverse lineages, reflecting variation in the mechanisms that govern entry into and exit from dormancy. Among bacteria in the phylum Actinomycetota, dormancy can be terminated by resuscitation-promoting factors (Rpf), exoenzymes that hydrolyze glycosidic bonds in the peptidoglycan of the bacterial cell wall. Although much of what is known about Rpf stems from studies on *Micrococcus luteus* and *Mycobacterium tuberculosis*, Rpf is present in many other Actinomycetota species and potentially in other phyla [21]. In this study, we characterized Rpf from *Micrococcus* KBS0714, a bacterial isolate from agricultural soil [26, 27]. We tested the effects of recombinant Rpf from this strain on a diverse set of dormant soil bacteria (Fig. 5). Patterns of resuscitation were related to the phylogeny of these bacteria and could be explained in part by conserved features of the cell envelope. Additionally, bacterial responses to Rpf were associated with functional traits, in particular, the moisture niche, which are critical for understanding persistence and resuscitation during soil dormancy. These findings expand our understanding of how Rpf may affect the diversity and functioning of terrestrial ecosystems.

### Kinetics and catalysis of Rpf

Rpf was initially characterized as a bacterial cytokine due to its small molecular size and its potent ability to resuscitate dormant cultures of *Micrococcus luteus* at picomolar concentrations (10^−12^ M) [17]. In contrast, we observed resuscitation at much higher Rpf concentrations (0.5–1.0 *µ*M), consistent with responses reported for other environmental isolates [24]. While some of these concentration-dependent effects may reflect the maximum rate of peptidoglycan hydrolysis (*V*_max_), it is also important to consider substrate affinity when characterizing enzyme kinetics. To this end, we determined that Rpf from *Micrococcus* KBS0714 has a low half-saturation constant (*K*_*m*_), indicating strong binding to peptidoglycan even at low substrate concentrations (Fig. 1). Although very few studies have systematically quantified the kinetics of Rpf, substantial variation in enzymatic activity have been observed, potentially reflecting differences in substrate (i.e., peptidoglycan) concentration and accessibility across environments.

Although Rpf typically stimulates the resuscitation of dormant bacteria at low concentrations, it can inhibit growth at higher concentrations. Similar to lysozyme, Rpf cleaves the *β*-1,4 linkage between N-acetylglucosamine (NAG) and N-acetylmuramic acid (NAM) in peptidoglycan. This breaking of this glycosidic bond initiates hydrolysis, facilitating cell wall remodeling necessary for growth and division. However, at elevated Rpf concentrations, excessive bond disruption or interference with regulatory processes involved in cell wall biosynthesis may compromise cell wall integrity. While inhibition at high Rpf concentrations has been reported [17, 21, 23, 28], we observed no evidence of this in our live cell assays (Fig.2). Instead, the growth of *Micrococcus* KBS0714 monotonically increased with Rpf concentration up to 6 *µ*M (Fig. 2). Taken together, our results suggest there is considerable variation among Actinomycetota strains in the concentration-dependent effects of Rpf on peptidoglycan hydrolysis and post-resuscitation growth.

### Genetic conservation of Rpf function

Resuscitation was strongly affected by site-directed mutations in highly conserved regions of the Rpf gene. The catalytic domain of Rpf found towards the N-terminal contains residues that are essential for muralytic activity (Fig. S3). Mutations in the glutamic acid residue (E54) significantly reduced or eliminated the ability of recombinant Rpf to resuscitate dormant *Micrococcus* KBS0714 (Fig. 3). These findings are consistent with other studies across various bacterial strains lending further support that the conserved glutamate residue is critical for peptidoglycan hydrolysis [19, 29].

Resuscitation was also affected by mutations in other regions of the protein. In the *rpf* sequence of *Micrococcus* KBS0714, we identified 21 repeating motifs in a lectin-encoding linker region between the catalytic domain and the LysM domain. In contrast, the *Micrococcus luteus* Flemming strain, which is a model for studying Rpf, contains only three repeats (Fig. S3) [17]. These repeats likely arose from tandem duplication due to replication slippage, which could have consequences for Rpf function. Lectins are involved in carbohydrate binding and can modify cell-substrate interactions. Deletion of these lectin repeats, along with the LysM domain, completely abolished resuscitation of dormant *Micrococcus* KBS0714, highlighting the functional significance of these repeating motifs. The additional lectins in Rpf may enhance peptidoglycan binding capacity and contribute to its high substrate affinity (Fig. 1). Thus, our findings lend support to view that there are conserved catalytic sites that are essential for resuscitation (e.g., E54), while also suggesting that structural variation (e.g., lectin repeats) is important for carbohydrate binding, which together may influence resuscitation in physically complex environments like soil.

### Population dynamics of resuscitated bacteria

One of the hallmarks of Rpf is that it reduces the lag time of dormant populations after waking up from dormancy [30]. The duration of lag time reflects the activity of cells and is influenced by the time required to carry out biosynthetic processes, such as the production of RNA and proteins that are necessary for growth and reproduction. Consistent with other reports, our experiments revealed that Rpf can significantly reduce lag time. Following 90 d of starvation, a one-time addition of recombinant Rpf allowed dormant *Micrococcus* KBS0714 to resuscitate one week earlier than control populations (Fig. 4). In an ecological context, accelerated growth responses could have major consequences for populations living in diverse communities, like soil, where there are rapid fluctuations in environmental conditions that affect microbial metabolism [31]. If Rpf enables dormant *Micrococcus* KBS0714 to resuscitate days ahead of other dormant bacteria, it could enter exponential phase earlier and persist, even in the presence of less responsive but potentially faster-growing competitors. This type of lottery effect may be further enhanced by reducing the stochastic nature of resuscitation [15, 32]. For instance, we observed that the among-population variation in both *µ*_max_ and lag time across replicate populations were significantly reduced (Fig. 4), indicating that Rpf may synchronize resuscitation-related processes and minimize random events that might otherwise disrupt regrowth.

### Rpf in a community context

At any given time, most microorganisms in soil are slow-growing or metabolically inactive [5]. The reactivation of dormant cells may be facilitated by Rpf. While Rpf is predominantly produced by bacteria within the phylum Actinomycetota, its target substrate, peptidoglycan, is a structural component of the cell wall that is found in nearly all bacteria. Additionally, the glycosidic bond cleaved by Rpf is chemically invariant, suggesting that a wide variety of dormant bacteria within the soil seed bank may be susceptible to resuscitation by this exoenzyme.

To better understand the breadth of resuscitation responses, we exposed a diverse collection of dormant soil bacteria to recombinant Rpf from *Micrococcus* KBS0714. Growth curve analysis revealed that bacteria do not respond uniformly to Rpf. Parameters such as yield and *µ*_max_ showed greater sensitivity to Rpf exposure, whereas lag time was less affected (Figs. 5, S4). Some of this variation could be explained by the evolutionary history of the strains used in our study. For example, there was a strong phylogenetic signal associated with the response of *µ*_max_ to Rpf. One likely factor contributing to this phylogenetic signal is the structural differences in the cell envelope, particularly the distribution and amount of peptidoglycan.

In Gram-positive bacteria, approximately 90% of the cell wall consists of peptidoglycan, forming a thick layer that is directly exposed to the external environment [33]. In contrast, Gram-negative bacteria have a thinner peptidoglycan layer, accounting for only 10% of the cell wall, which is protected between the inner and outer membranes [33]. Our results indicate that Rpf from *Micrococcus* KBS0714 had a stronger positive effect on yield and *µ*_max_ in Gram-negative bacteria compared to Gram-positive bacteria. Because there is more peptidoglycan exposed to the extracellular environment, we expected that Rpf would have a stronger effect on Gram-positive bacteria. In contrast, we found that Gram-negative bacteria were more responsive to Rpf, suggesting that other mechanisms besides direct contact with substrate (peptidoglycan) are important for the resuscitation of dormant bacteria. In any case, cell envelope is a coarse-grained trait that may be useful for making predictions about Rpf-mediated resuscitation in complex communities.

The response of diverse bacterial strains to Rpf may be influenced by other functional traits. In soils, microbial activity is often governed by moisture. When water availability is low — due to precipitation, soil texture, or evapotranspiration — microbial activity typically drops, causing cells to enter a dormant state. When water availability suddenly increases, for example, following a rain event, microorganisms often exhibit pulses of metabolic activity as they exit dormancy [31]. Understanding how microorganisms respond to these fluctuations, and how this controls transitions into and out of dormancy, has important implications for the diversity and function of soil microbiomes. We found that the response of dormant bacteria to Rpf was correlated with moisture niche parameters [26]. Specifically, dry-adapted strains with a narrow niche breadth were less responsive to Rpf, while wet-adapted strains with a broader moisture niche breadth were more responsive to Rpf (Figs. 6). Another potential adaptation to low moisture conditions is biofilm production [34]. Our results revealed a positive correlation between the Rpf effect on lag time and biofilm production, suggesting that a smaller investment into biofilms may allow strains to resuscitate more readily. This could indicate a trade-off or reflect differences in enzyme activity and diffusion within the exopolymeric matrix.

While Rpf serves as a potentially widespread mechanism for terminating bacterial dormancy, many other factors contribute to microbial seed-bank dynamics. Although much remains to be elucidated, it is known that Rpf acts synergistically with Rpf-interacting protein A (RipA), an endopeptidase that cleaves stem peptides involved in peptidoglycan cross-linking. RipA localizes at the septa of growing cells, facilitating cell wall digestion and releasing small muropeptides, which may serve as signaling molecules, thereby imparting specificity to the resuscitation process [35, 36]. Likewise, Rpf-mediated resuscitation operates alongside, and may even interact with, other dormancy strategies such as endosporulation and persister cell formation, imparting additional dimensions to microbial survival [5]. Consequently, individual cells may exist in shallow or deep dormancy states of dormancy, all while possessing distinct histories, traits, and reserves that influence their longevity and responsiveness to internal and external cues. Collectively, these processes create structure and memory that can give rise to rich and emergent phenomena [8]. Despite this complexity, the findings presented here contribute to our understanding of dormancy regulation in diverse microbial habitats, such as soils, with implications for understanding indirect effects, feedbacks, and the stability and functioning of ecosystems [25].

## 4 Methods

### 4.1 Strains and culturing

We used *Micrococcus* KBS0714, an actinomycetotal strain that was isolated from agricultural soil [26]. It is very closely related to *M. luteus* NCTC 2662, an isolate that has been used to study Rpf [17]. The genomic characteristics and physiology of KBS0714 have been described in detail elsewhere [26, 27]. For routine culturing, we maintained KBS0714 on R2B liquid medium at 25 C on an orbital shaker (150 rpm). Unless otherwise stated, we induced dormancy by allowing cells to enter stationary phase and then keeping them on a shaker table (150 rpm) for 30 d followed by static (non-shaken) conditions for an additional 60 d. Prior to using these dormant bacteria for resuscitation experiments, we washed the cells by pelleting and resuspending them five times in phosphate buffered saline (PBS, pH = 7.0) to remove residual medium.

### 4.2 Recombinant protein expression

We amplified and cloned the *rpf* gene of *Micrococcus* KBS0714 into an *Escherichia coli* expression host to produce recombinant Rpf. As described in greater detail elsewhere [25], we extracted genomic DNA from KBS0714 using a Microbial DNA isolation kit (MoBio). We then amplified the open reading frame of the *rpf* gene using two primers, Upper-F 5’ GCC CAT ATG GCC ACC GTG GAC ACC TG 3’ and Lower-R 5’ GGG GAT CCG GTC AGG CGT CTC AGG 3’, which incorporated the EcoRI restriction sites NdeI (forward primer) and BamHI (reverse primer) [17, 37]. The PCR conditions are as follows: initial: 95 C for 5 min, 30 cycles of 95 C for 30 s, 55 C for 30 s, 72 C for 1 min, and final extension at 72 C for 7 min. The PCR product was confirmed using gel electrophoresis and Sanger sequencing [25]. We performed an NdeI/BamHI restriction digest on the 850 bp *rpf* gene amplicon and ligated the restriction product into a pET15b expression vector with an N-terminus polyhistidine-tag (pET15B-His_6_-rpf). We then transformed the recombinant expression plasmid into the *E. coli* Origami BL21 (DE3) expression host (*E. coli* pET15b-His_6_-rpf).

To overexpress Rpf, we grew *E. coli* pET15b-His_6_-rpf in 1 L of Lysogney Broth (LB) and induced protein expression at an OD600 of *∼*0.6 with isopropyl *β*-D-1-thiogalactopyranoside (IPTG) (final concentration 0.1 mM). We then lysed the cells via sonication, which was followed by centrifugation and filtration (0.45 *µ*m). With this preparation, we purified the recombinant Rpf protein with the N-terminus polyhistidine-tag via Ni-NTA Purification (Invitrogen) using a 10 mL gravity-fed column with a 2 mL resin bed. The protein was run through the column twice and rinsed with 3X volume of wash Ni-Buffer A (300 mM NaCl, 50 mM Tris-HCl, 5 mM imidazole) followed by elution with 125 mM imidazole elution buffer in 1 mL fractions. Rpf was further purified via buffer exchange using 10 mL Zeba Spin Desalting Columns (Thermo Fisher) with protein buffer (20 mM Tris-HCl, 100 mM NaCl) according to manufacturer’s instructions followed by 0.2 *µ*m syringe filtration. Overexpression and protein purity were confirmed by SDS-PAGE and Western blotting (Fig. S1). Protein concentrations were determined with Bradford assays (Invitrogen).

### 4.3 Enzyme kinetics

We quantified the enzymatic activity of recombinant Rpf with the EnzChek Lysozyme Assay Kit (Molecular Probes). The assay estimates muralytic activity via the degradation of fluorescein-infused peptidoglycan. We incubated Rpf (1 mg/mL, 34 µM) over a range of substrate concentrations (0 - 30 µg/L) at 22 C for 24 h. As a control, we deactivated the enzyme via incubation at 90 C for 1 h. Fluorescence was then measured with a BioTek Synergy H1 plate reader using excitation//emission wavelengths of 485//530 nm. We quantified the enzyme kinetics by fitting the data with a Michaelis-Menten function using maximum likelihood procedures [38]:

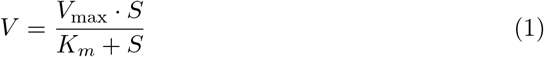

where *V* is the rate of peptidoglycan hydrolysis, *V*_max_ is the maximum rate of peptidoglycan hydrolysis, *K*_*m*_ is the half-saturation constant, and *S* is the substrate concentration.

### 4.4 Concentration dependence of Rpf

We characterized the growth of *Micrococcus* KBS0714 in response to a concentration gradient of recombinant Rpf. After growing *Micrococcus* KBS0714 to late stationary phase (30 d), dormant cells were washed 5X in PBS and then diluted 10,000-fold before being spread-plated (100 µL) in triplicate onto R2A agar plates containing 0 - 6 textmuM Rpf. The plates were then incubated at 25 C for 96 h. Colony biomass was determined via image analysis with Adobe Photoshop. We fit the data to a Monod growth equation, which is identical in form to the Michaelis-Menten function used for characterizing enzyme kinetics for Rpf (see eq. 1).

### 4.5 Population dynamics

We used a growth-curve assay to quantify the resuscitation dynamics of *Micrococcus* KBS0714 in response to a one-time addition of recombinant Rpf. After washing 5X in PBS, we inoculated an equal number of dormant cells into a set of 250 mL side-arm flasks containing 35 mL of lactate minimal medium with 1% L-lithium lactate [39]. In quadruplicate (n = 4), the flask were amended with recombinant Rpf (0.5 µM final concentration) or a protein buffer as a control. We measured biomass over time as optical density (OD600) using a Biophotometer (Eppendorf) while the flasks were incubated on a shaker (150 rpm) at 25 C. We fit the resulting data to a modified Gompertz equation [40] using maximum likelihood procedures:

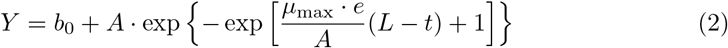

where *µ*_max_ is the maximum growth rate (h^−1^), *λ* is lag time (h), and *A* is the carrying capacity or biomass yield (OD600), and *b*_0_ is the intercept. We tested for the effect of the Rpf treatment on the growth parameters using using *t*-tests. We also quantified the within-replicate coefficient of variation (CV) of the growth-curve parameters to test whether Rpf affects the synchrony of resuscitation.

### 4.6 Rpf mutations

We created mutations in the *rpf* gene of *Micrococcus* KBS0714 to better understand the mechanisms of Rpf activity. First, we mutated the invariant glutamate residue (E54) of Rpf using QuikChange Site-Directed Mutagenesis Kit (Stratagene) following the manufacturer’s instructions. Conditions and reagents for the mutagenesis are described in Supplementary Tables S1 (primers sets), Table S2 (PCR master mix), and Table S3 (PCR conditions). We methylated the parental DNA by incubating with 1 µL of DpnI enzyme at 37 C for 1 h. With the resulting mutated DNA, we transformed competent *E. coli* (Invitrogen) using 5 µL of PCR product to generate a population of transformants that grew on LB plates containing 100 µg/mL of ampicillin. We grew these cells for *∼*16 h in 5 mL of LB with the appropriate antibiotics on a shaker (150 rpm) at 37 C. We then extracted plasmids from the cells using QIAprep Spin Miniprep Kit following the manufacturer’s instructions. We confirmed the success of the mutagenesis via Sanger sequencing [41]. Second, we truncated a variable linker and LysM domain of the native *rpf* gene using PCR amplification, cloning, transformation, and Rpf expression as described in the previous sections. Specifically, two primers, rpf1-F 5’ACC GCG ACC GTG CAG CGC TAG GAT CCG 3’ and trncrpf-R 5’CGG ATC CTA GCG CTG CAC GGT CGC GGT 3’ were designed to PCR amplify the N-terminus lysozyme-like region omitting the variable linker and LysM Domain. The PCR resulted in a*∼*375 bp amplicon as determined by gel electrophoresis and Sanger sequencing. We then overexpressed and purified the recombinant protein for each of the mutant sequences following the procedures described above. Dormant cells were washed 5X in PBS prior to being incubated in 13 mm test tubes with 2 mL of R2B broth containing 2.5 µM Rpf on a shaker table (150 rpm) at 25 C for 96 h. Growth was measured as biomass (OD600) on replicate (n = 4) test tubes using a Biophotometer (Eppendorf). We compared growth of *Micrococcus* KBS0714 exposed to Rpf (native vs. mutated) and the negative control using one-way ANOVA and Tukey’s HSD.

### 4.7 Diverse bacterial responses to Rpf

We evaluated the specificity of Rpf by exposing different strains of dormant bacteria to the recombinant protein that was expressed from the *rpf* gene in *Micrococcus* KBS0714. We used 12 soil strains that are part of a genomically and physiologically well-characterized culture collection [26, 42]. Gram-negative bacteria included *Pedobacter* KBS0701, *Azospirillium* KBS0705, *Pseudomonas* KB0710, *Janthinobacterium* KBS0711, and *Variorvorax* KBS0712. Gram-positive bacteria included *Arthrobacter* KBS0702, *Arthrobacter* KBS0703, *Mycobacterium* KBS0706, *Microccous* KBS0714, *Curtobacterium* KBS0715, *Rhodococcus* KBS0724, and *Bacillus* (KBS0812). Each strain was grown in batch culture containing R2A broth medium on a shaker (150 rpm) at 25 C. After being maintained in stationary phase for 30 d, we washed the dormant cells 5X in PBS and dispensed them into replicate wells (n = 4) of Corning Costar flat-bottom 48-well culture plates (Thermo Fisher) with 1 mL R2A broth that contained 0.5 µmol/L recombinant protein (+Rpf) or an equal volume of 100 mM Tris-HCl protein buffer (-Rpf). We then measured growth as optical density (OD600) using a BioTek plate reader (Thermo Fisher) at 25 C under constant shaking (150 rpm). With the resulting data, we estimated growth curve parameters by fitting a modified Gompertz model (see eq. 2). With these parameters, we calculated the Rpf effect size for each strain using Cohen’s 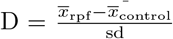where 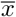 is the mean value of a growth parameter and sd is the pooled standard deviation across treatment levels.

We evaluated the effect of Rpf with respect to the evolutionary history of the bacterial strains. With existing 16S rRNA gene sequences [26, 42], we created a maximum likelihood tree with RAxML that used a General Time Reversible (GTR) model with the Gamma model of rate heterogeneity. We mapped the Rpf effect size for growth curve parameters (*µ*_max_, lag time, and biomass yield) onto the phylogenetic tree using the *ape* package in R [43]. We then tested for phylogenetic signal of the Rpf effect size using Blomberg’s K and Pagel’s *λ* with the *phytools* package in R [44].

We evaluated the effect of Rpf in relation to functional traits that were previously measured on the bacterial strains [26]. These traits included biofilm production, motility, and microaerotolerance. In addition, we characterized the moisture niche of each strain by measuring rates of respiration along a gradient of soil water potential (MPa). With a modified Gaussian function, we extracted variable that describe the microbial water niche, including the minimum moisture, the optimal moisture, and the breadth around the moisture optimum [26]. We then used simple linear regression to describe relationships between functional traits and the Rpf effect size.

