## Supplementary Information for "Resuscitation-promoting factor (Rpf) terminates dormancy among diverse soil bacteria"

1

| Primers | Sequence |
|---|---|
| 5' Rpf E54A | cctcgccgagtgcGCGtccagcggcacc |
| 3' Rpf E54A | ggtgccgctggaCGCgcactcggcgagg |
| 5' Rpf E54Q | cctcgccgagtgcCAGtccagcggcacc |
| 3' Rpf E54Q | ggtgccgctggaCTGgcactcggcgagg |
| 5' Rpf E54K | cctcgccgagtgcAAGtccagcggcacc |
| 3' Rpf E54K | ggtgccgctggaCTTgcactcggcgagg |

| Reagent | Volume ($\mu$L) |
|---|---|
| $H_2O$ | 35 |
| 10X Phusion HF buffer | 5 |
| 100% DMSO | 4 |
| 25 mM dNTPs | 1 |
| 50 mM $MgCl_2$ | 1 |
| 5' Rpf mutagenesis primer, 4 $\mu$M | 1 |
| 3' Rpf mutagenesis primer, 4 $\mu$M | 1 |
| Rpf template plasmid, 15 ng/$\mu$L | 1 |
| Phusion enzyme | 1 |
| **Total** | **50** |

**Table 1. Experimental conditions for Rpf mutagenesis.** Upper: primer sets and sequences used for site-directed mutagenesis of the conserved glutamate residue in *rpf* from *Micrococcus* KBS0714. Upper-cased bases indicate introduced mutations in the sequence. Lower: reagents for master mix used in PCR for site-directed mutagenesis

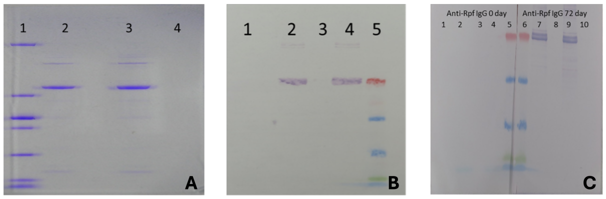


**Fig 1. Rpf over-expression gels.** **A**. SDS-PAGE of pET15b Rpf at 900 µg/mL confirming protein purity extracted from NI-NA column extraction. First lane is a low-range protein marker (100, 30, 25, 20, 15, 10, 5, 3.4 kDa), second lane is 5 µL of Rpf, third lane is empty, fourth lane is 10 µL of Rpf. **B** Western blot of 5 and 10 µL of pET15b recombinant Rpf at 900 µg/mL concentration bound with anti-Histidine (1:50000) primary IgG in lane 2 and 4. Lane 1 and 3 are empty. Lane 5 is a low-range western marker (red/40, blue/15, green/10, blue/2.6, blue/1.7kDa). **C** Western blot of 5 and 10 µL of pET15b Rpf protein at 900 µg/mL concentrations bound with 72 day Rpf-specific primary IgG antibodies (1:50000) in lanes 7 and 9. This is contrasted with the 0 day antibodies which did not bind to Rpf protein in lane 2 and 4. Lanes 1, 3, 8, and 10 are empty. Lanes 5 and 6 are low-range western marker (red/40, blue/15, green/10, blue/2.6, blue/1.7kDa)

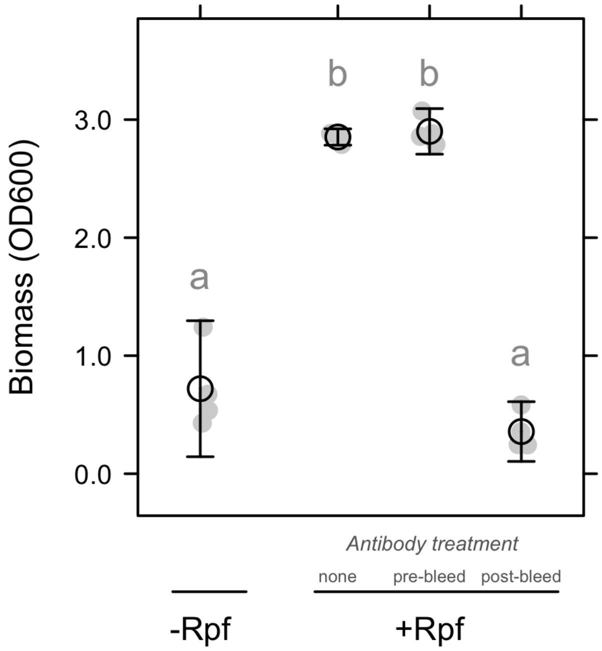


**Fig 2. Assessment of negative controls.** We performed an experiment where we measured the regrowth of dormant *Micrococcus* KBS0714 in R2A broth medium with and without Rpf. Final biomass was estimated at 120 h of a growth curve assay. In the negative control (-Rpf), we only added protein buffer. In the +Rpf treatments, cells were exposed to 1.57 µM of recombinant Rpf. Within the +Rpf treatment, there was an antibody treatment where cells were exposed to no antibody (”none”) or 600 µg/mL concentration of either 0-day Rpf-antibodies (”pre-bleed”) 28-d Rpf-antibodies (”post-bleed”). Antibody binding to Rpf active site of muralytic activity is sufficient for abolishing resuscitation effects on KBS0714 biomass growth to levels comparable to protein buffer control.

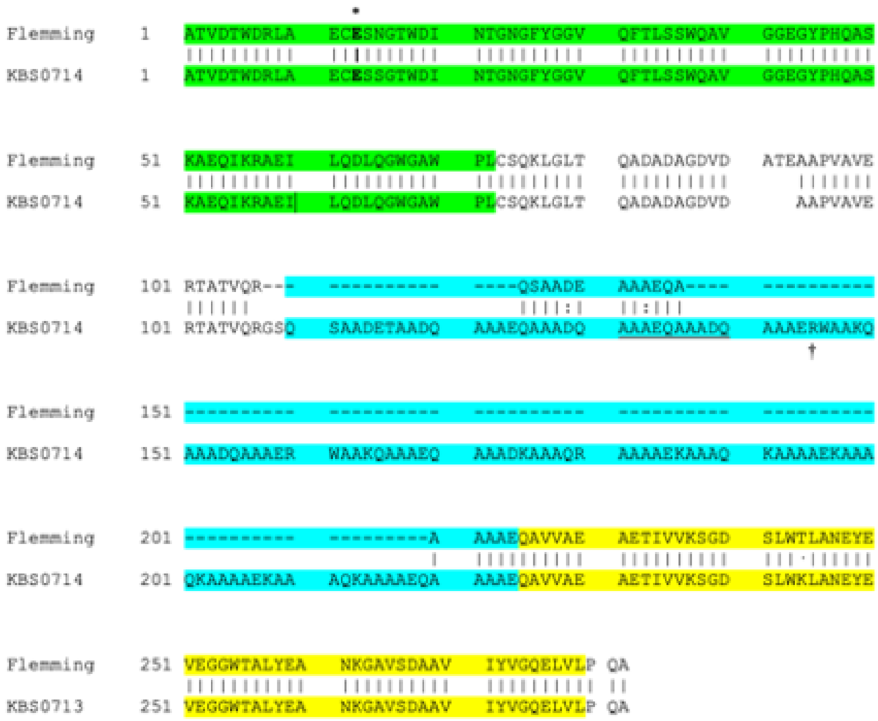


**Fig 3. Amino acid sequences of Rpf.** Amino acid sequences for resuscitation promoting factors (Rpf) from *Micrococcus luteus* (Flemming) and a soil isolate used in this study, *Micrococcus* KBS7014. Starting at residue 42, there is a conserved glutamic acid (E) residue at position 54 (*), which is within the lysozyme-like domain highlighted in green. We made recombinant Rpf with KBS014 strain that had site-directed substitutions at the conserved site to evaluate effects on resuscitation (See Fig. 3). The lectin-rich linker region, the length of which is greatly reduced in the Flemming strain (-), is highlighted in blue while the LysM domain is highlighted in yellow. We made a recombinant Rpf with a KBS0714 strain that truncated a significant fraction of the lectin-rich linker region and the LysM domain starting at residue 145 (†). Other symbols follow UniProt convention where solid vertical lines represent fully conserved residues, colons (:) represent residues with strongly similar properties, and periods (.) represent residues with weakly similar properties.

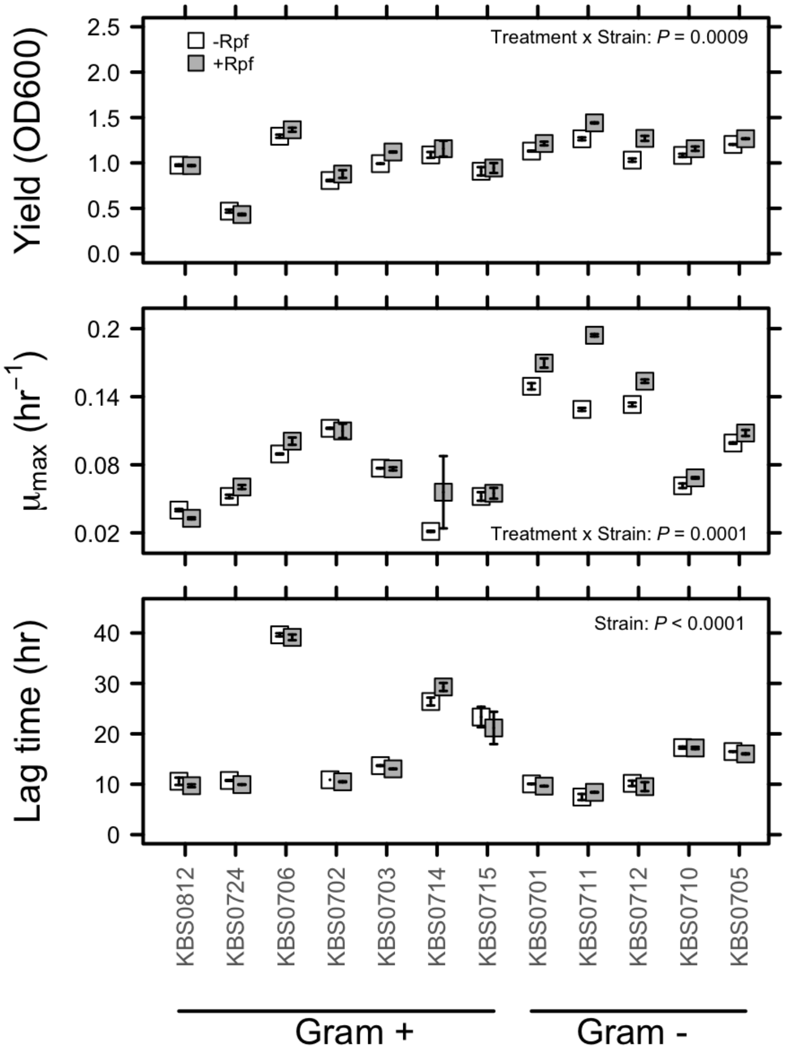


**Fig 4. Growth parameters.** Growth parameters for different soil bacterial when exposed to recombinant protein (+Rpf) or a negative control (-Rpf)
